# GRANAR, a new computational tool to better understand the functional importance of root anatomy

**DOI:** 10.1101/645036

**Authors:** Adrien Heymans, Valentin Couvreur, Therese LaRue, Ana Paez-Garcia, Guillaume Lobet

## Abstract

Root hydraulic conductivity is an important determinant of plant water uptake capacity. In particular, the root radial conductivity is often thought to be a limiting factor along the water pathways between the soil and the leaf. The root radial conductivity is itself defined by cell scale hydraulic properties and anatomical features. However, quantifying the influence of anatomical features on the radial conductivity remains challenging due to complex, and time-consuming, experimental procedures.

We present a new computation tool, the Generator of Root ANAtomy in R (GRANAR) that can be used to rapidly generate digital versions of root anatomical networks. GRANAR uses a limited set of root anatomical parameters, easily acquired with existing image analysis tools. The generated anatomical network can then be used in combination with hydraulic models to estimate the corresponding hydraulic properties.

We used GRANAR to re-analyse large maize (Zea mays) anatomical datasets from the literature. Our model was successful at creating virtual anatomies for each experimental observation. We also used GRANAR to generate anatomies not observed experimentally, over wider ranges of anatomical parameters. The generated anatomies were then used to estimate the corresponding radial conductivities with the hydraulic model MECHA. This enabled us to quantify the effect of individual anatomical features on the root radial conductivity. In particular, our simulations highlight the large importance of the width of the stele and the cortex.

GRANAR is an open-source project available here: http://granar.github.io

**One-Sentence summary:** Generator of Root ANAtomy in R (GRANAR) is a new open-source computational tool that can be used to rapidly generate digital versions of root anatomical networks.

## Introduction

Root water uptake is influenced by a multi-scale combination of structural and hydraulic properties ^1,2^. At the root system scale, the overall root architecture defines the potential uptake sites within the soil domain. Hydraulic properties of individual roots (radial and axial conductivities) further constrain these uptake sites, thus establishing a global hydraulic architecture of the plant ^3^. Changes in these local hydraulic properties, which have a global influence, are thought to be an important target in breeding programs for drought resistant crops ^4^.

At a finer scale, on the organ level, radial and axial conductivities can also be defined by hydraulic and structural properties. On the hydraulic side, the radial conductivity (k_r_) of a root segment is influenced by the expression and location of water channels ^5^, the formation of hydrophobic barriers ^6^, and the conductivity of plasmodesmata ^7^. On the structural side, the k_r_ is known to be influenced by anatomical features, such as the number of cell layers in the cortex ^8^, the size of cortex cells ^9^, or the presence of aerenchyma ^10^. Generally speaking, a plant controls its hydraulics in short and medium-term ^11^, and its structural features over longer developmental timelines ^12^. As such, root anatomy defines the baseline for the root radial hydraulic properties.

Root anatomical traits have been shown to be diverse within species such as maize^13,14^ or cassava ^15^, between species^16^, and in different biomes ^17^. However, to our knowledge, an assessment of the functional implication of anatomical features in water uptake properties has never been done in a systematic way. Indeed, the in vivo quantification of radial and axial conductivities is usually performed using a root pressure probe ^18^, which is experimentally restrictive, because it is hard to perform on soil-grown roots and is generally seen as a low-throughput procedure. This experimental restriction has hindered the investigation of the relation between root hydraulic properties and anatomical features.

Recently the development of a modelling tool of transverse water flow in roots from the sub-cellular scale, MECHA, opened the way to the systematic analysis of the influence of anatomical and cell hydraulic properties on k_r_ ^7^. A remaining limitation of the approach is that it requires explicit anatomical networks as input. Such networks can be obtained by digitizing cross section images with semi-automated tools, such as CellSet ^19^. However, that procedure is time consuming for large datasets. Therefore, there is a need for a high throughput method of acquisition of root anatomical networks ^2^.

Here, we present a new computational tool, the Generator of Root ANAtomy in R (GRANAR) that can generate digital versions of root anatomical networks. GRANAR uses easily accessible anatomical features as input. These tissue-scale parameters can be acquired from open-access image analysis softwares and root cross section images. In combination with the model MECHA, GRANAR opens up the new possibility to systematically analyse the impact of root anatomy on hydraulic conductivity in a high-throughput fashion. GRANAR has been validated using published datasets of maize anatomical data and hydraulic measurements. GRANAR is an open-source project available at https://granar.github.io/.

## Results

### Overview of the Generator of Root ANAtomy in R

The GRANAR model generates cell networks of root cross sections from a small set of root anatomical features (Table 1). The root anatomical features can typically be gathered by using software image analysis, such as ImageJ ^20^, or with more automated software, such as RootScan ^21^, PHIV-RootCell ^22^ or RootAnalyzer ^23^. Briefly, the anatomy generation process is based on the placing of cell layers around the center of the root. The position and size of each cell and cell layer is a function of the cell type radius and user-defined randomness. As a result of this randomness, each simulation produces a slightly different anatomy, even with the same input parameter set. The thickness of a specific tissue is therefore a function of its cells’ size and its number of layers. Specific vascular tissue patterns are created depending on whether the user chooses to simulate a monocot or a dicot anatomy (fig. 1). Independent of the species, in figure 1, GRANAR was able to reproduce cross section anatomies with high similitude (RE: 3.4%, R^2^: 0.99) over the total area of the cross section between the experimentally obtained cross section and the one obtain with GRANAR.

**Table 1:** List of parameters used by the model to generate root cross section.

| Tissues | Parameter |  |  |
| --- | --- | --- | --- |
| **Stelar parenchyma** | Cell diameter | Diameter of the stele tissue not including the pericycle |  |
| **Pericycle** | Cell diameter | Number of layers |  |
| **Endodermis** | Cell diameter | Number of layers |  |
| **Cortex** | Cell diameter | Number of layers |  |
| **Exodermis** | Cell diameter | Number of layers |  |
| **Epidermis** | Cell diameter | Number of layers |  |
| **Xylem** | Max cell diameter | Number of xylem poles | Ratio (# *protoxylem* vs # *metaxylem*) |
| **Aerenchyma** | Proportion | Number of radial cavities |  |

**Figure 1:**
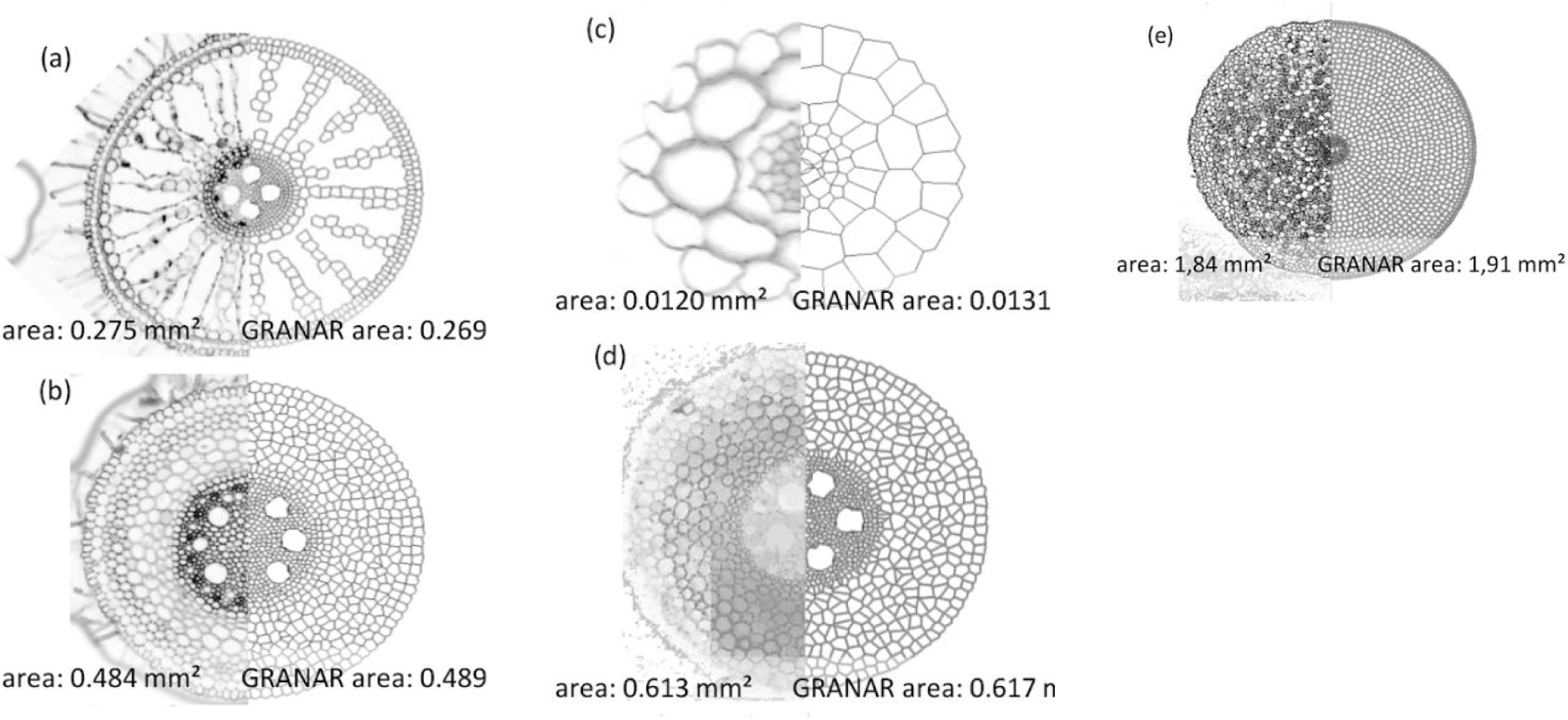
Visual comparison between root cross section images of different plants and their simulation with GRANAR. **(a)** Rice (*Oryza sativa*) adventitious root grown hydroponically. Collected 10 cm above the root tip. **(b)** Wheat (*Triticum aestivum*) cv. Savannah mature adventitious root grown in soil. Collected 5 cm from stem. **(c)** *Arabidopsis thaliana* primary root grown on an agar plate. Collected 5 cm from the root tip. Initial image cross section comes from ^24^, figure 4. **(d)** Maize (*Zea mays. B73 line*) crown root grown aeroponically. Collected 5 cm from the root tip. Initial image cross section comes from ^25^. **(e)** Mature Ranunculus root ^26^. The model parameters used for those simulation were gathered with ImageJ and are available at: https://github.com/granar/granar_examples. The area information below each image correspond to the total area of the cross sections.

Once the root anatomy is created, the cell network can be saved in a file in eXtended Markup Language (XML). The file in XML can serve as the geometry input for the MECHA model. As such, both tools can be linked, and the GRANAR-MECHA model can be used to estimate hydraulic properties (e.g. k_r_) of any type of root anatomy. The format of the XML created by GRANAR are identical to the ones created by CellSet, which also generates a fully digitize root anatomical network. Using the same format for both tools makes them modular and facilitates the intercomparison of their outputs (fig. 2).

**Figure 2:**
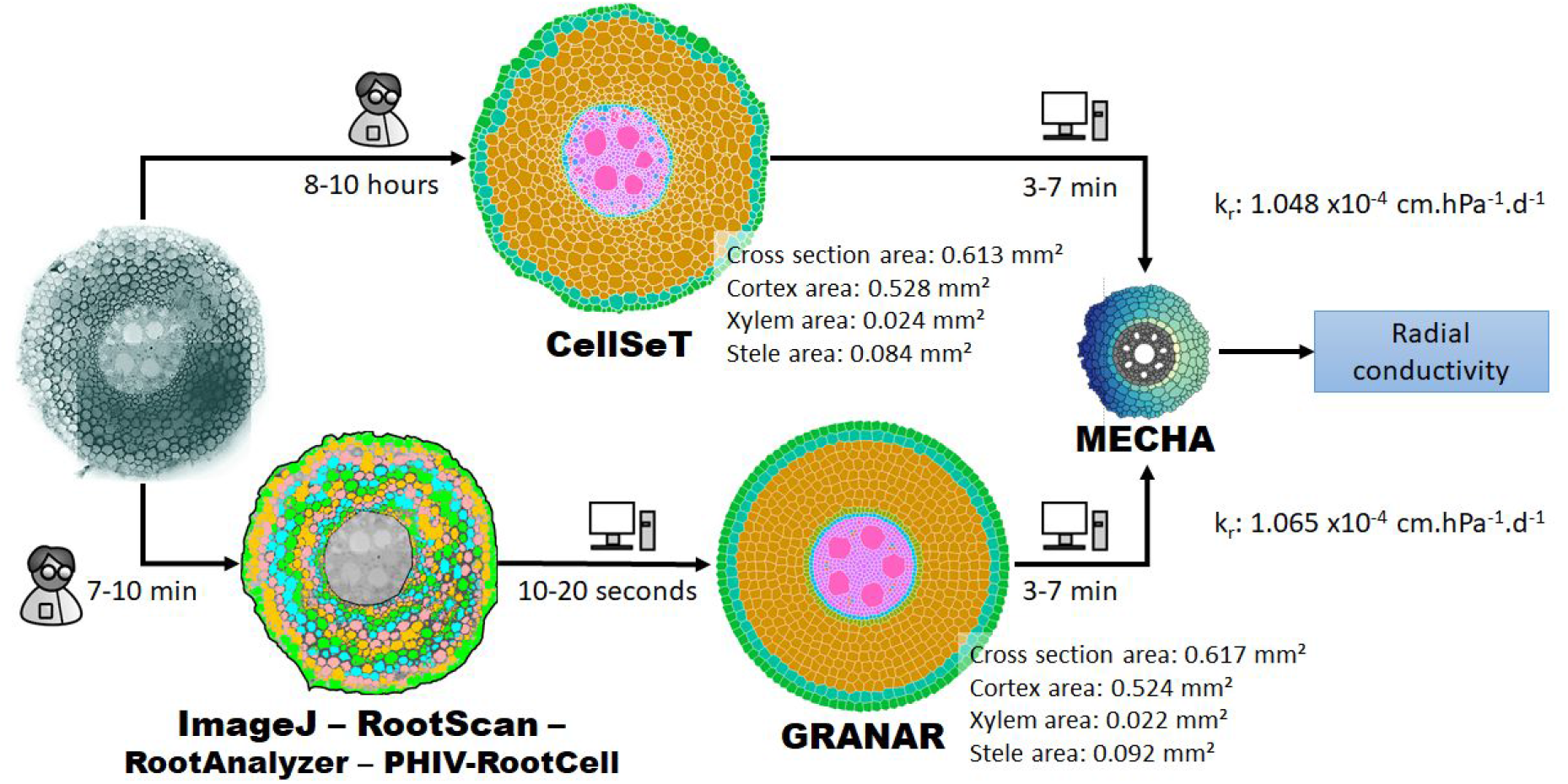
Overview of the methods to shortcut the full digitalization with CellSet of a root anatomy and access an estimation of single-root scale hydraulic properties (*k_r_* = radial hydraulic conductivity). The input parameter of GRANAR in the figure were gathered with ImageJ. The *k_r_* values were calculated using the MECHA model assuming an endodermal casparian strip as the only apoplastic barrier and uniform cell-scale hydraulic properties. The person- or computational-time required for the procedure are detailed next to each arrow.

The time (both computational and labor time) required to access the level of network detail with CellSet in figure 2 is around 10 hours in cross sections with thousands of cells, such as maize (image blending and correction (30 minutes) + automated segmentation (3 seconds) + iterative corrections of the segmentation and labelling (~10 hours)). In comparison, generating a complete network anatomy only takes 10 minutes (Root anatomical features (7 minutes RootScan/image or 10 minutes ImageJ/image) + GRANAR/cross section (15 seconds)). However, when we compare the k_r_ obtained by MECHA using either Cellset or GRANAR anatomy as input, the difference is less than 2 percent (Figure 2).

### GRANAR generates accurately stereotypic maize root anatomies from large anatomical datasets

We validated the output of GRANAR for maize (*Zea mays*), whose root anatomy is well documented in the literature. We used the experimental data from three independent datasets showing large phenotypic diversity of maize anatomical features. The three root anatomical datasets were from Burton et al. (2013a) (195 landraces from a diversity panel), from Chimungu et al. (2014b) (38 maize genotypes with large variance of cortex features), and from Gao et al. (2015)^27^ (1 cultivar with 3 different root types and a large proportion of aerenchyma in all the 3 root types). These different datasets contain tissue-scale anatomical data that could be used as input for GRANAR, such as the size of the stele or the number of cortex layers (details about how these data were compiled to create the GRANAR input files are presented in the Material and Methods section). Using the 239 data points, we generated 717 maize root cross sections. Each data point was used to generate 3 replicates of each root, taking advantage of the randomness in the positions of the individual cells.

Once all the anatomies were created, we compared organ-scale metrics (that were not directly used as input to create the anatomies) between the experimental and simulated data (fig. 3). We can observe that GRANAR is able to accurately represent the total areas and sizes of the different tissues of the root (e.g. fig 3a: total cross section area, fig 3b: total cortex area, fig. 3c: xylem area, fig. 3d: aerenchyma area). For each variable analyzed, the r-squared value between the simulated and experimental data were above 0.95. The relative error of GRANAR is below 5% for the total cross section area, the total cortical area and the xylem area, and less than 15% for the aerenchyma area. For each variable analysed, no significant differences were found between experimental and simulated data (P > 0.05).

**Figure 3:**
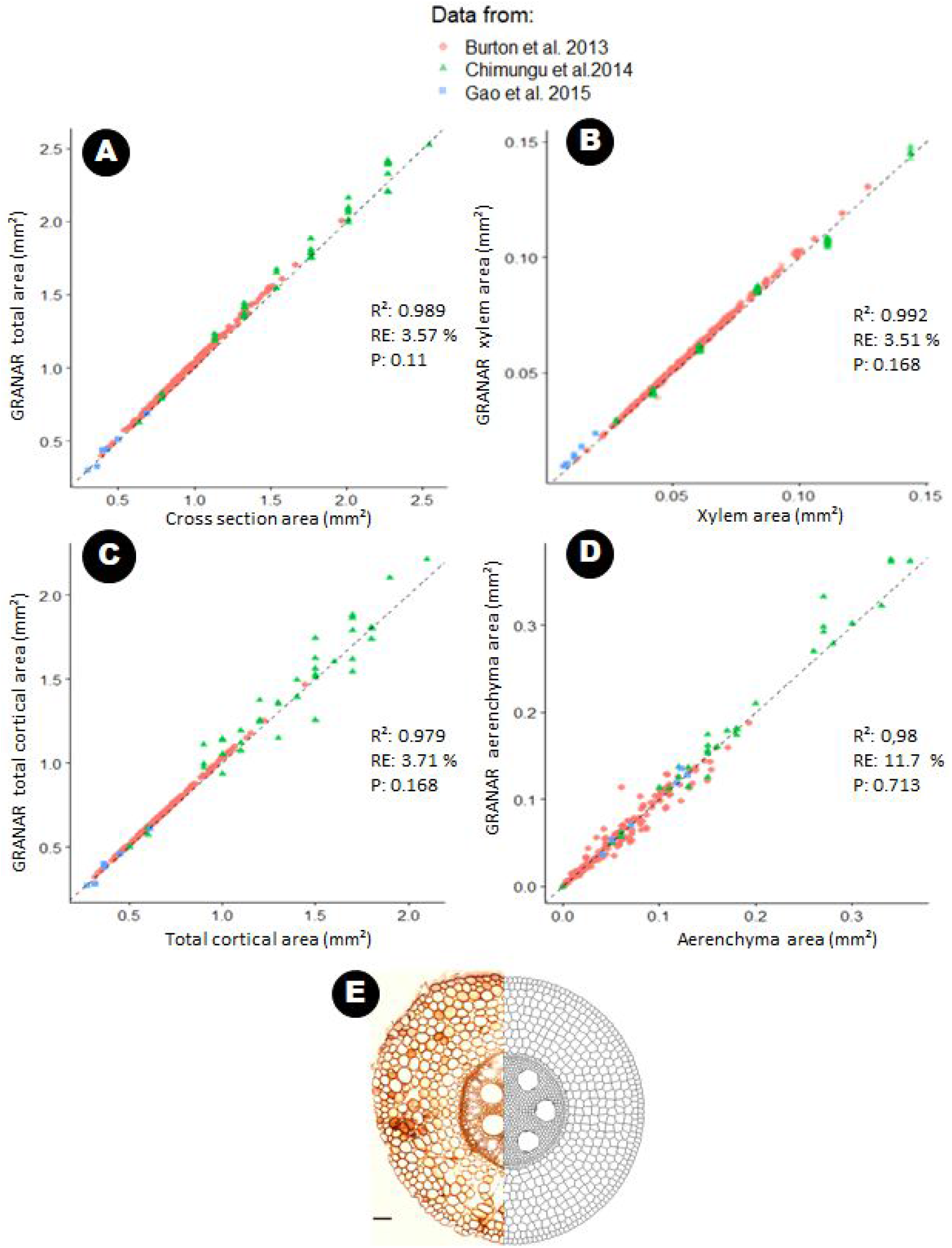
Comparison between simulated and experimental anatomies for 4 anatomical features. A: Total cross section area. B: Cortical area. C: Xylem area. D: Aerenchyma area. The different colors and shapes represent the different datasets. RE = relative error; R^2^: R-squared, P: p-value for the F test. E: illustration depicting one initial image (Chimungu et al. 2014) and the generated anatomy side-by-side. Bar = 100μm

### GRANAR-MECHA enables the estimation of radial conductivity from simulated anatomical data

By design, the anatomical output of GRANAR can be used as an input for the MECHA model ^7^. We were therefore able to estimate the k_r_ for each of the experimental data points, assuming constant cell-scale hydraulic properties (see Material and Methods for details). The simulated k_r_ (across the different root cross sections) vary between 21.1 and 8.1 x 10^−8^ m.MPa^−1^.s^−1^ assuming an endodermal Casparian strip as the only apoplastic barrier.

Using the results of these simulations, we were able to attribute the main changes of k_r_ to three parameters using a linear regression model (table 2). The simulated k_r_ for those data points can than be estimated using the following equation:

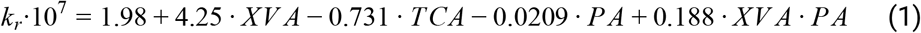

where k_r_ is the simulated radial conductivity (m MPa^−1^ s^−1^), XVA is the xylem area (mm^2^), TCA is the total cortical area (mm^2^), and PA is the aerenchyma proportion (dimensionless [0,100]).

**Table 2:**
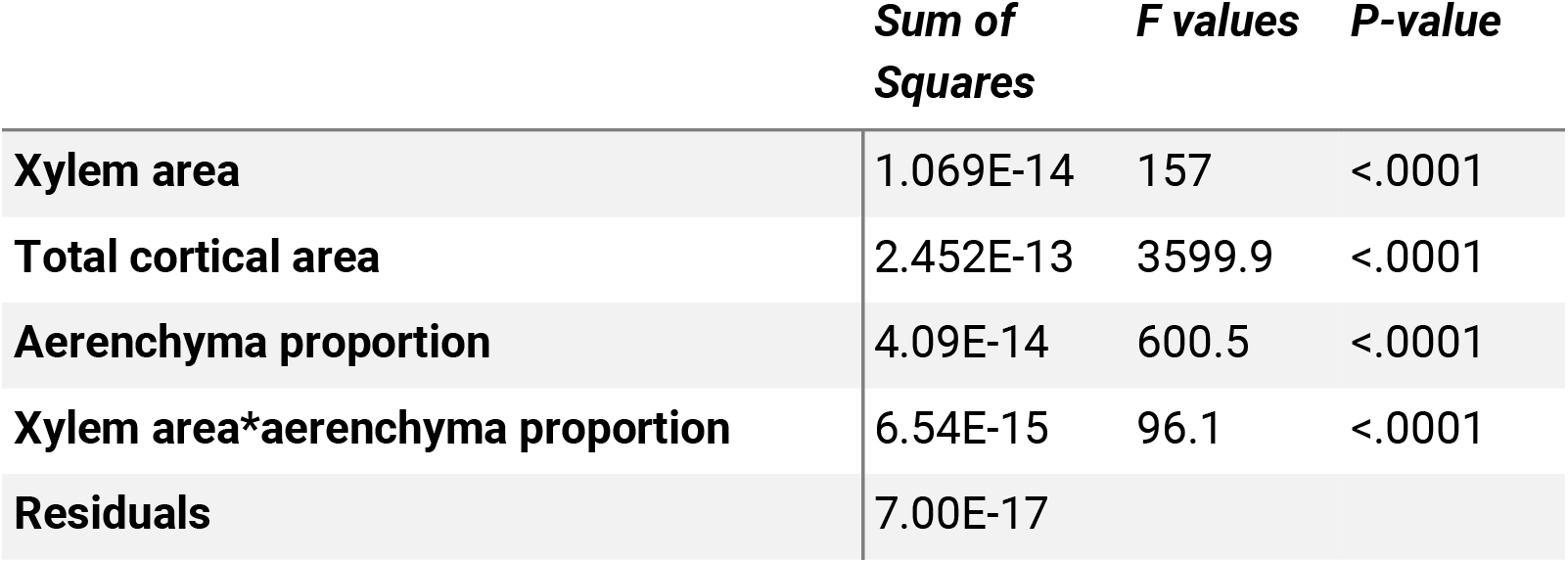
Summary of the ANOVA for the effect of the xylem and the total cortical area combined with the aerenchyma proportion on the k_r_ when the only apoplastic barrier is an endodermal Casparian strip.

|  | **Sum of Squares** | **F values** | **P-value** |
| --- | --- | --- | --- |
| **Xylem area** | 1.069E-14 | 157 | <.0001 |
| **Total cortical area** | 2.452E-13 | 3599.9 | <.0001 |
| **Aerenchyma proportion** | 4.09E-14 | 600.5 | <.0001 |
| **Xylem area\*aerenchyma proportion** | 6.54E-15 | 96.1 | <.0001 |
| **Residuals** | 7.00E-17 |  |  |

The most substantial correlation factor to the simulated k_r_ was found for the cortex width (corr: −0.84; P < 0.05, table 3). As a general rule, as the cortex width increases, the k_r_ decreases (fig. 4a). In addition, for a given cortex width, as the xylem area increases, the k_r_ increases (fig. 4a). The ratio between the stele area and the total area of the cross section is also highly correlated to the k_r_ (corr: 0.79, P < 0.05, table 3). In this regards, as the ratio between the stele and the cross section area increases, the k_r_ increases (fig. 4b). Additionally, for a given ratio between the stele and the cross section area, as the xylem area decreases, the k_r_ increases (fig. 4b). Moreover, in the analysed dataset, an increase in aerenchyma proportion seems to only slightly decrease the simulated k_r_ (fig. 4c). The simulated k_r_ of our study and the ones measured in the literature for maize were in the same range (fig. 5).

**Table 3:**
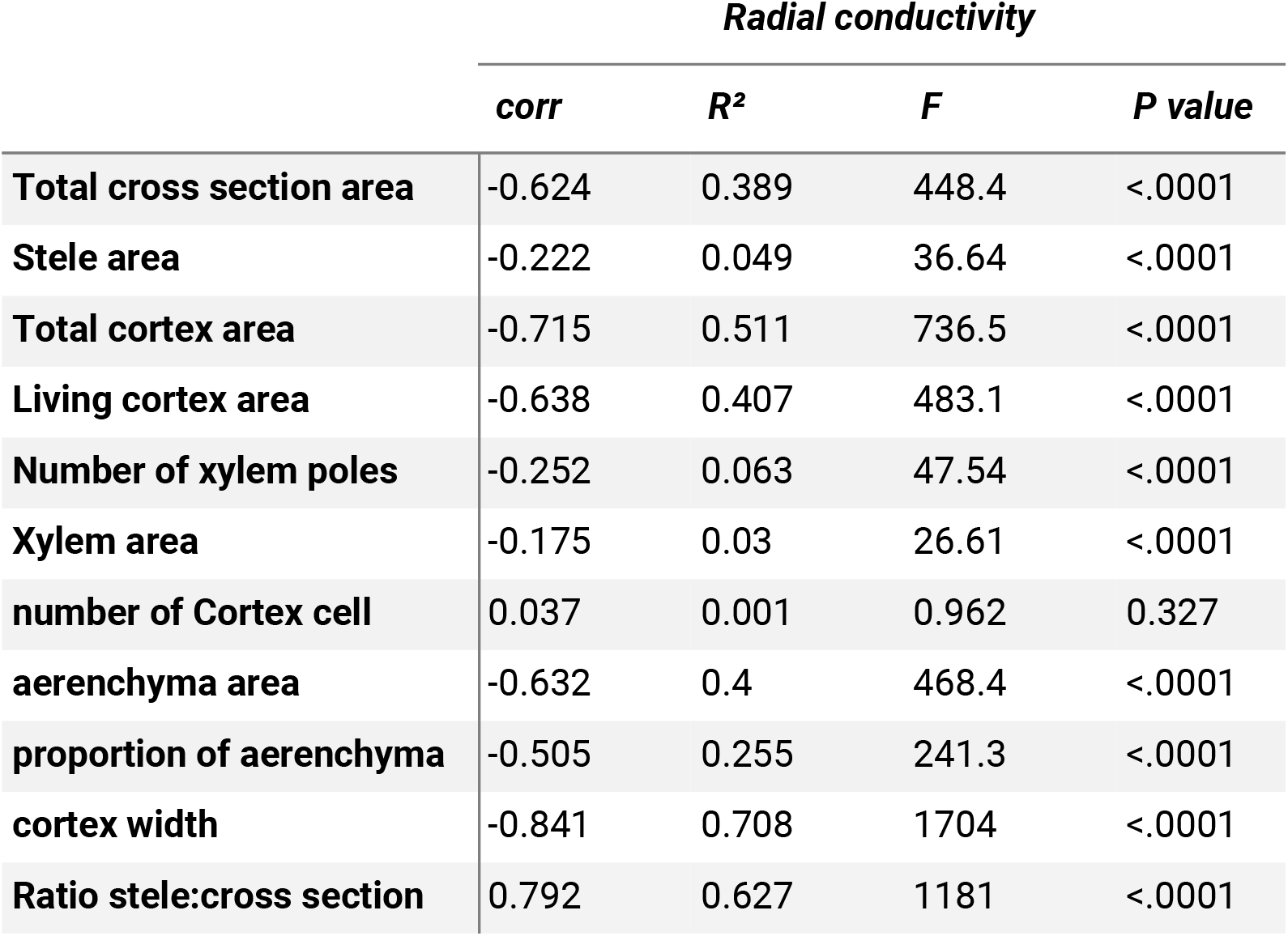
Summary of the ANOVA for the effect of one-by-one anatomical feature on the simulated k_r_ assuming an endodermal casparian strip as the only apoplastic barrier. Each line shows results from a single one-way ANOVA. corr = correlation coefficient of Pearson. R^2^ = R squared value

**Table 4:**
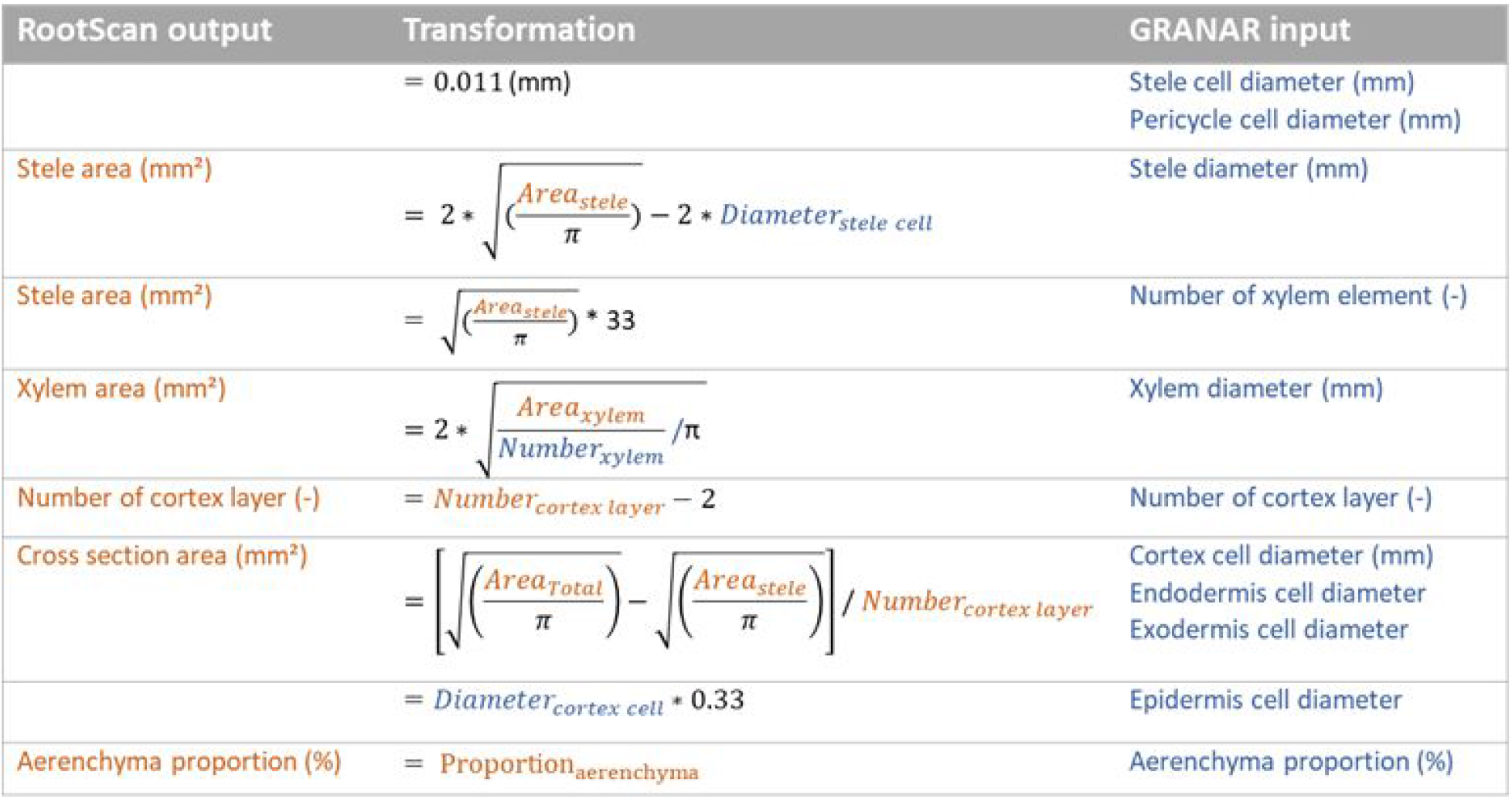
Overview of the methodology to transform RootScan data into GRANAR parameter.

**Figure 4:**
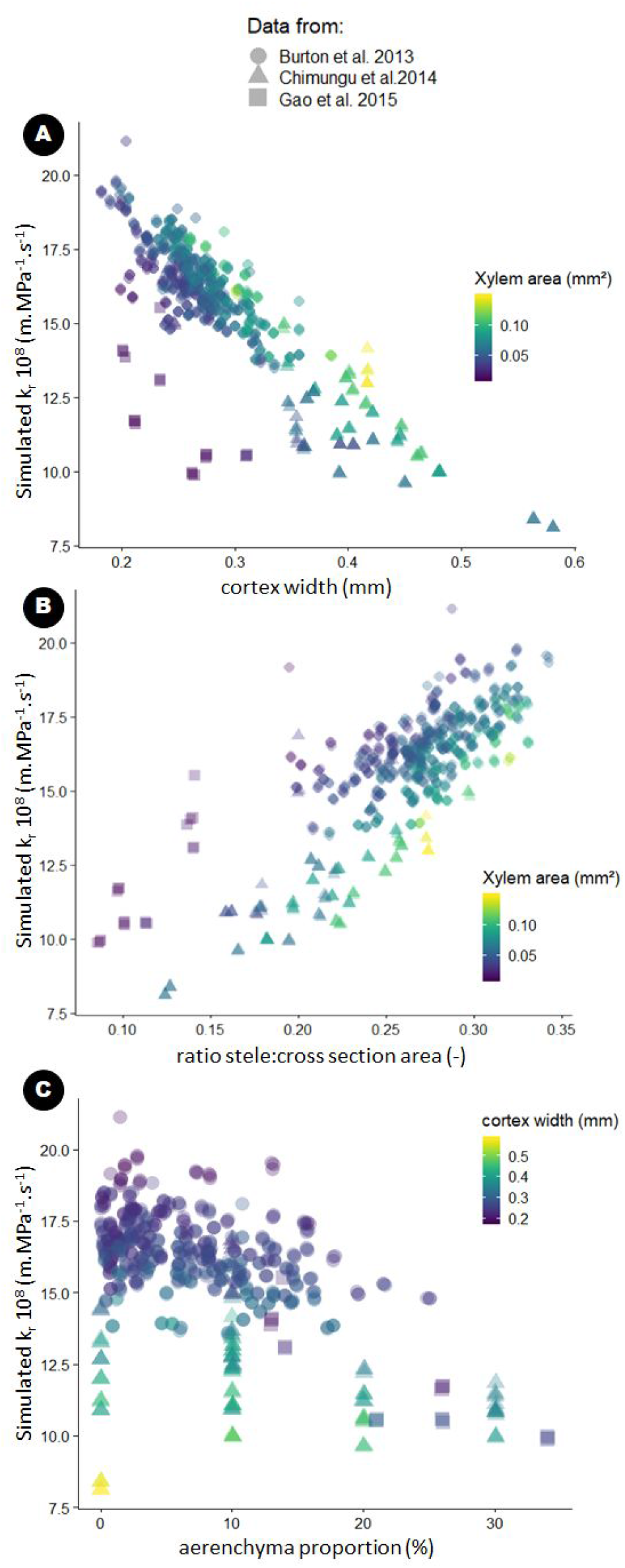
Relation between simulated k_r_ and anatomical features for each dataset (symbol). Scenario with endodermal casparian strip. **(A)** Effect of the cortex width on the simulated k_r_. Colors represent the xylem area. **(B)** Effect of the ratio between the stele area and the total cross section area. Colors represent the xylem area. **(C)** Effect of aerenchyma presence on the simulated k_r_. Colors represent the cortex width.

**Figure 5:**
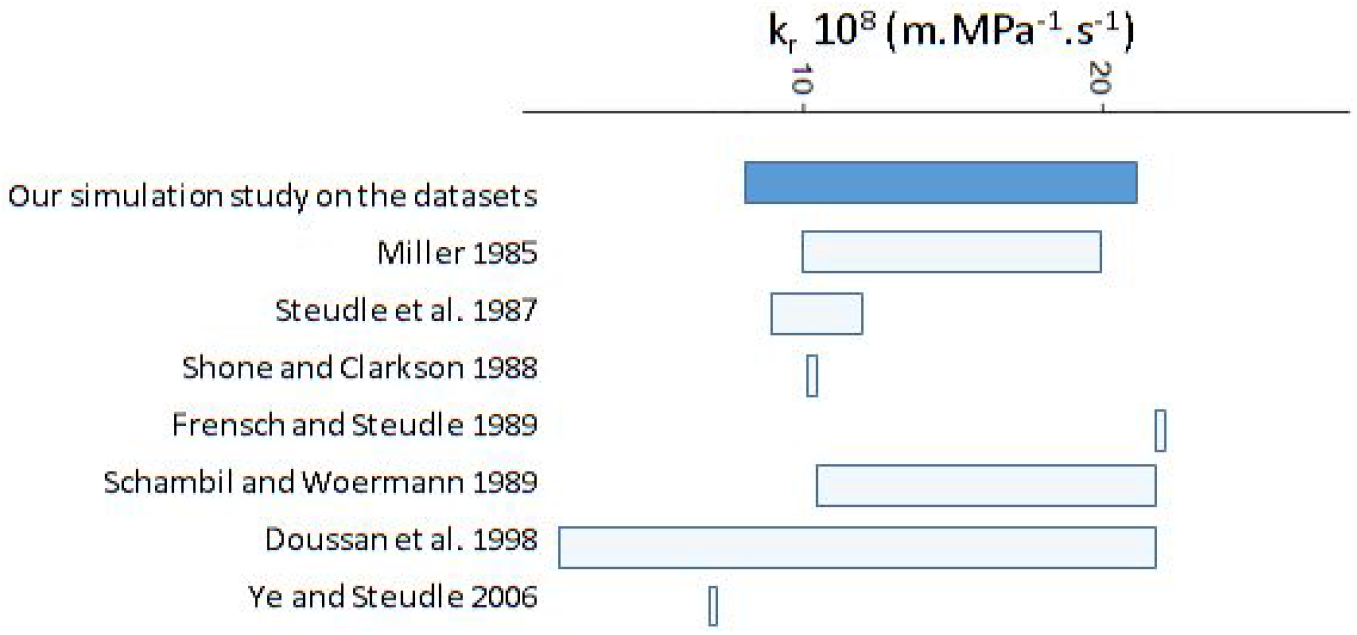
Comparison between the simulated k_r_ of our study and the ones measured in the literature for maize ^28–33^.

### GRANAR-MECHA enables sensitivity analysis of anatomical features on root radial conductivity

In the previous section, we used the GRANAR-MECHA model to study the theoretical effect of experimental root anatomical features on the root k_r_. The advantage of a modelling approach is that each anatomical feature can be changed independently from the others, which is often impossible experimentally. The capability to change anatomical features independently allows for a more complete exploration of the theoretical phenotypic spectrum.

Here, we chose to vary the proportion of aerenchyma, the size of the cortex, the size of the stele as well as the number and area of xylem poles. For each trait combination, we computed the k_r_ for three different maturity levels (fig. 6). The maturity levels account for three approximate distances from the root tip corresponding to different levels of apoplastic barrier development. Based on ^34^ and ^35^ we assumed the following apoplastic barrier types for the different distances: 1 = presence of endodermal Casparian strip (5 cm); 2 = presence of suberized endodermis (25 cm); and 3 = presence of suberized endodermis and exodermal Casparian strip (30 cm).

**Figure 6:**
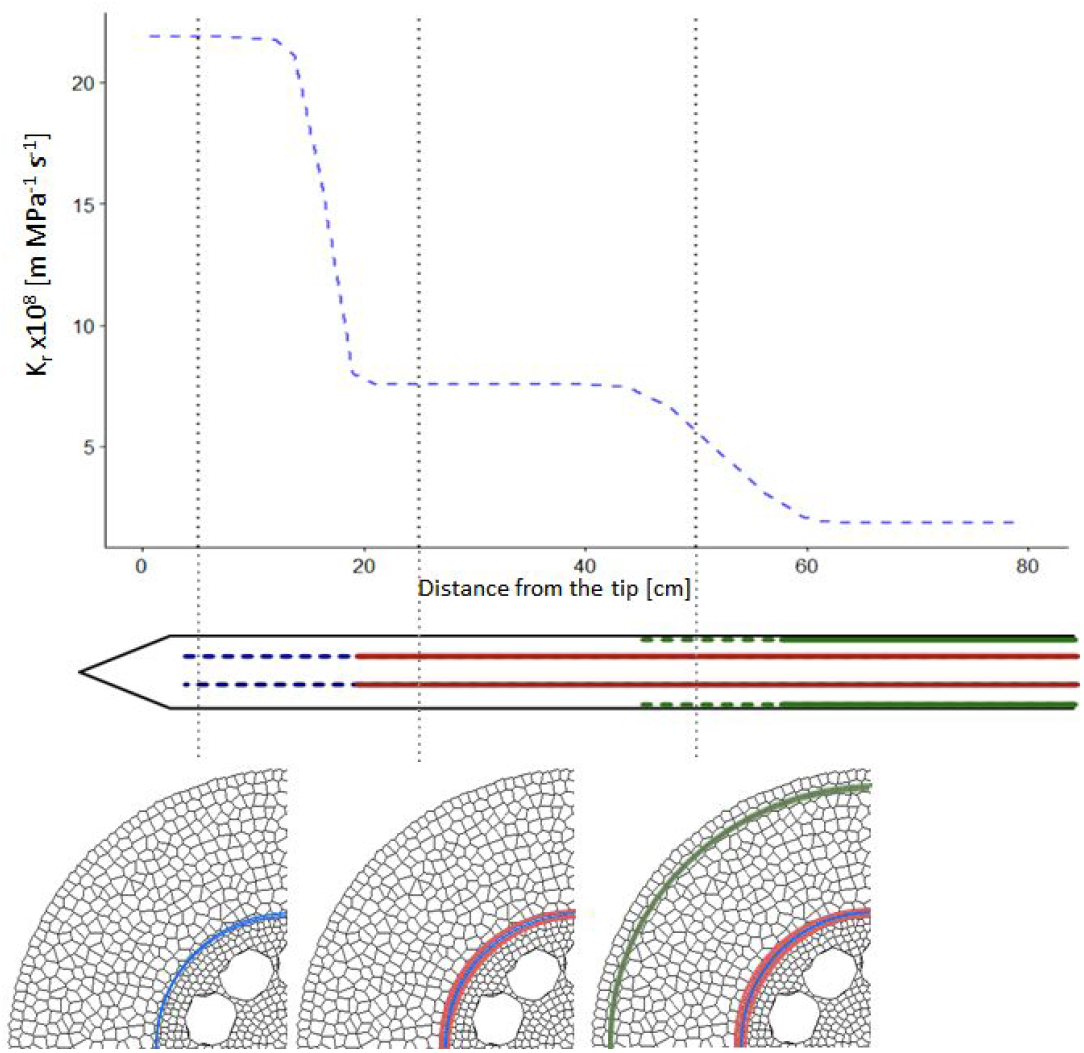
Estimated evolution of the k_r_ in main axes of maize root system derived from Doussan et al. (1998). Below, the illustration depict our assumption of the evolution of main apoplastic barriers based on Enstone and Peterson (2005) study. The different lines represent different level of apoplastic barriers. 1: Endodermal Casparian strip (dashed blue); 2: Endodermal suberization (red); 3: Endodermis full suberization and exodermal Casparian strip (dashed green).

The first anatomical feature that we chose to vary was the proportion of aerenchyma, from zero to fifty percent of the cortex area (fig. 7). The range observed in the experimental datasets spanned zero to thirty five percent (see previous section). An increase in aerenchyma induced a decrease in the simulated k_r_. The effect was less important as the endodermis started to suberize. However, under the maturity level 3, with a fully suberized endodermis and an exodermal Casparian strip, the relative k_r_ decrease is stronger than the one observed at the maturity level 2, with a suberized endodermis (fig. 7). The stronger relative decrease can be explained by the reduced amount of cells after the exodermis layer which allow the water to pass from a cell-to-cell to an apoplastic pathway whereas there is no shifting of preferential pathway of water next to the exodermis at maturity level 2.

**Figure 7:**
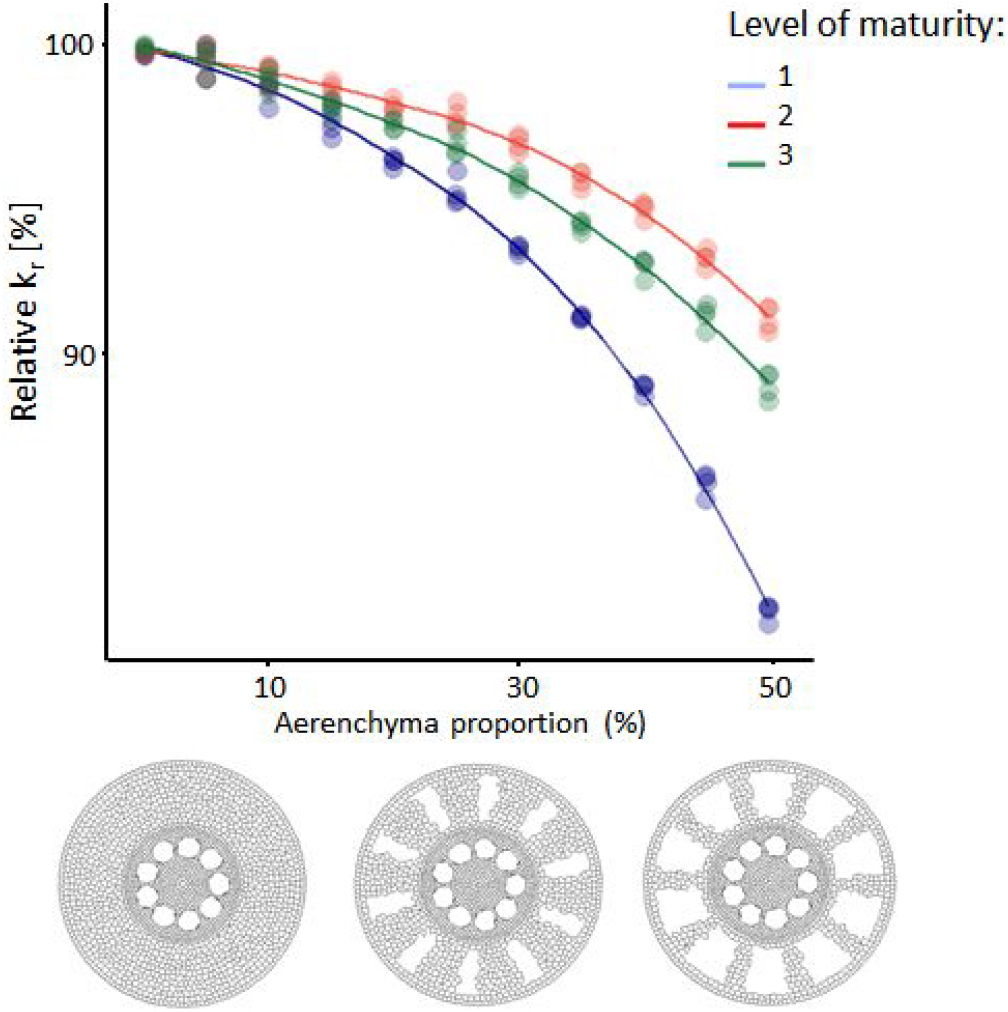
Influence of the proportion of aerenchyma on the relative k_r_ for simulated monocot anatomies. The colors represent different levels of hydrophobic barrier development. 1: Endodermal Casparian strip (blue); 2: Endodermal suberization (red); 3: Endodermis full suberization and exodermal Casparian strip (green). The cross sections at the bottom of the figure show, from left to right, examples of roots with 0%, 25% and 50% of aerenchyma.

To observe the influence of the cortex features on the k_r_, a range of cross sections were simulated with increasing cortex cell diameter and number of cortex layers (fig. 8). We observed that an increase of all components induces a decrease in the simulated k_r_ (fig. 8A-C-D). Moreover, the cortex width (isolines in fig. 8A) accounts for the coupled effect of the cell diameter and the number of layers on the simulated k_r_. This feature neatly captures the decreasing trend of k_r_ with increasing cortex width for the global simulated dataset included in fig. 8B (unlike figs. 8C and 8D displaying a single transect across the parametric space each).

**Figure 8:**
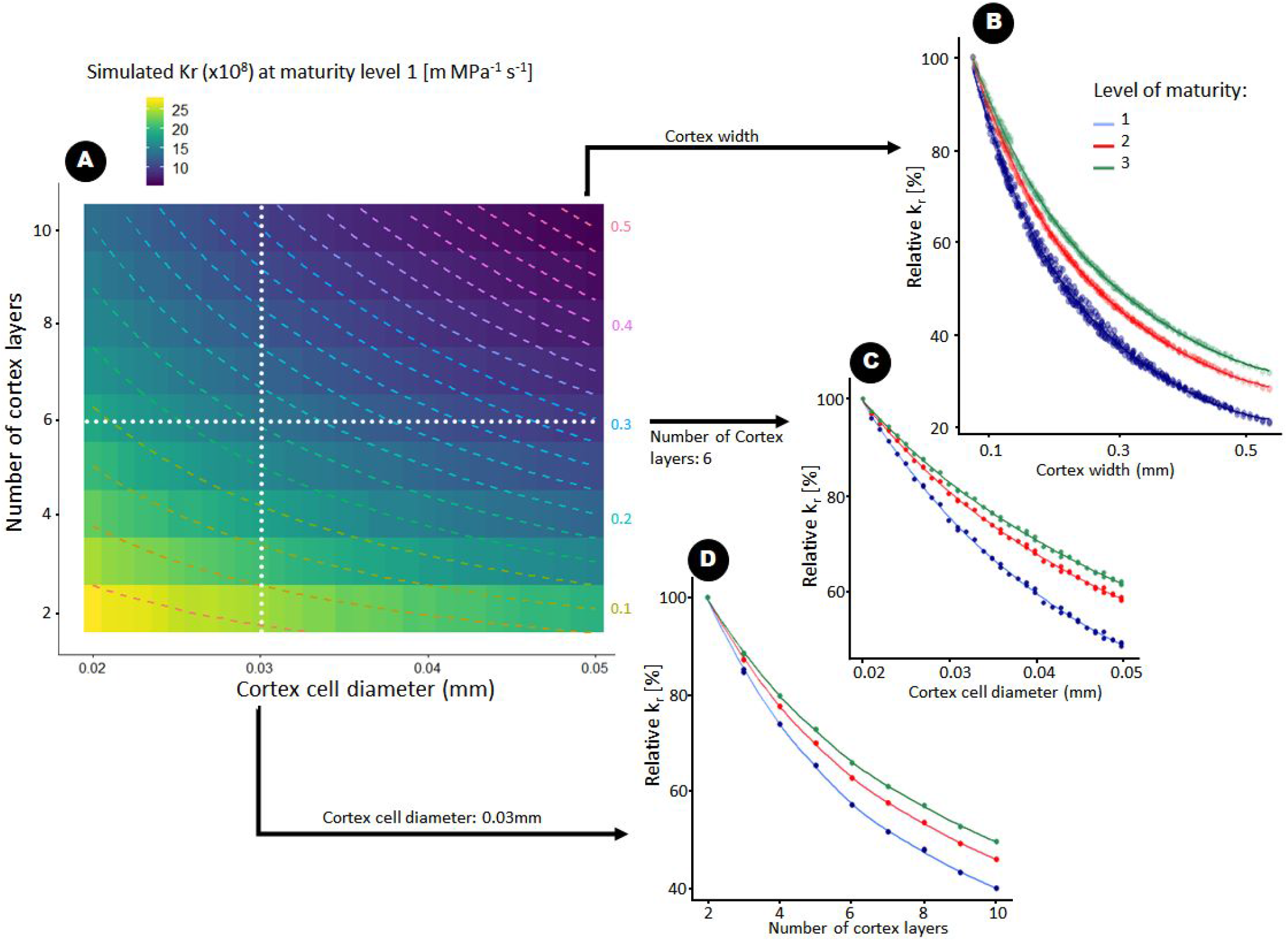
Influence of cortex anatomy (number of cortex layers and cell size) on k_r_ for simulated monocot anatomies. **(A)** Overview of k_r_ values across the parametric space in case of endodermal Casparian strip as the only apoplastic barrier. The colored dashed lines represent the cortex width isoline (mm) and the dotted lines are a visual representation of the fixed parameters for the two subplots (*C-D*). **(B)** Evolution of the relative k_r_ as the cortex width increases. All data points of the sub figure (*A*) are represented in blue. **(C)** Evolution of the relative k_r_ as the cortex cell diameter increases for a number of cortex layers fixed to six. **(D)** Evolution of the relative k_r_ as the number of cortex layers increases for a cortex cell diameter set to 0.03 mm. **(B-C-D)** The colors represent different levels of hydrophobic barrier development. 1: Endodermal Casparian strip (blue); 2: Endodermal suberization (red); 3: Endodermis full suberization and exodermal Casparian strip (green).

The relative influence of each cortex feature is greater at maturity level 1 than for the other maturity level tested. The reason is that the relative influence of the other apoplastic barriers to water flow is less encounter. Moreover, under maturity level 3, the cortex features have the lowest relative influence on the conductivity in comparison to the other ones.

In figure 9, simulations where the cortex width was kept constant and we increased the stele diameter. Increasing the stele area leads to an increase of the simulated k_r_ (fig. 9). The influence of the stele area is highest in the scenario with the maturity level 2.

**Figure 9:**
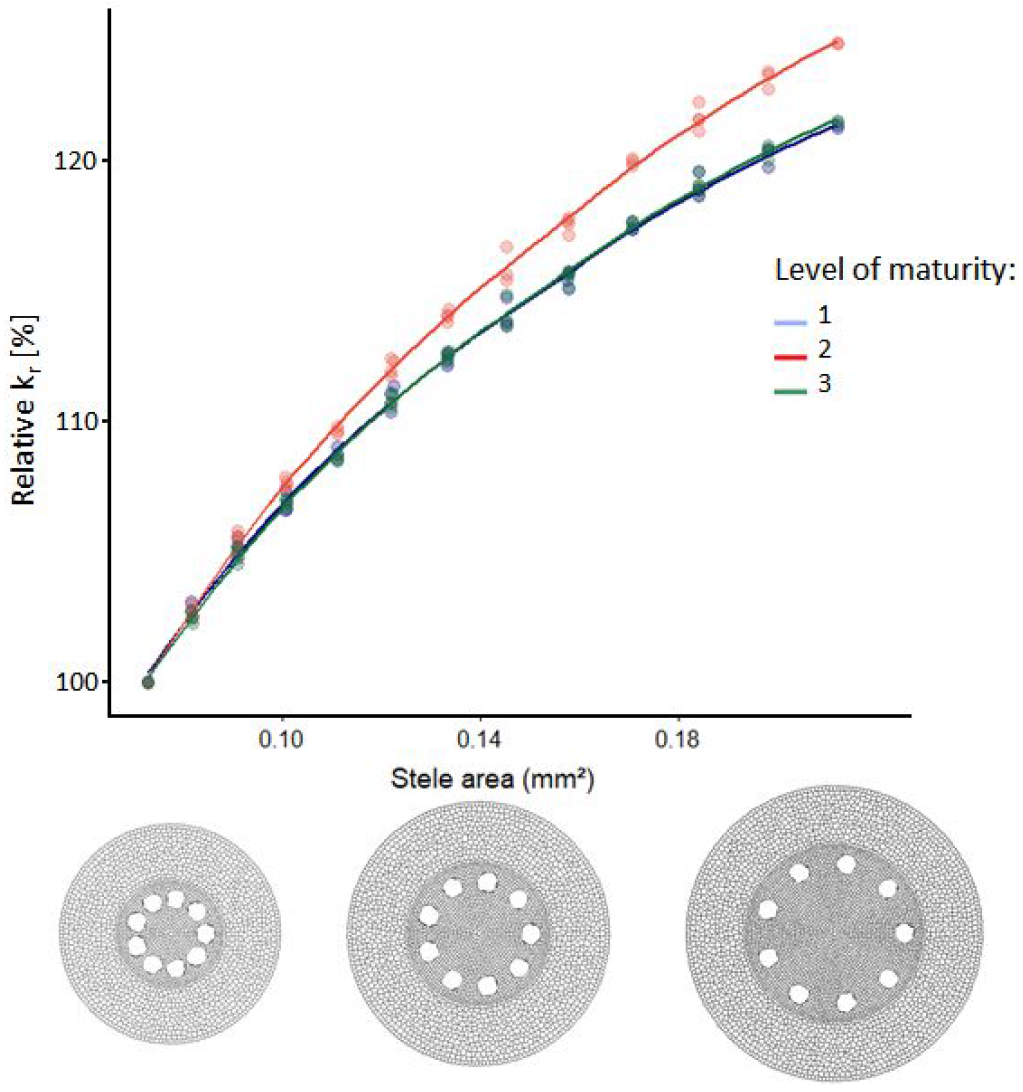
Influence of the stele area on the relative k_r_ for simulated monocot anatomies. The colors represent different levels of hydrophobic barrier development. 1: Endodermal Casparian strip (blue); 2: Endodermal suberization (red); 3: Endodermis suberization and exodermal Casparian strip (green). The cross sections at the bottom of the figure show, from left to right, examples of roots with 0.3, 0.4 and 0.5 mm stele diameter.

Figure 10 shows the influence of increased xylem area on the k_r_. To increase the xylem area, the number of the xylem poles was increased as well as the size of each pole. As a general rule, having more xylem poles increases the simulated k_r_. However, the k_r_ reached a plateau when the xylem nearly covered the whole stele perimeter (fig. 10A). At maturity level 1, the relative influence of the increase of xylem number is the greatest compared other maturity levels.

**Figure 10:**
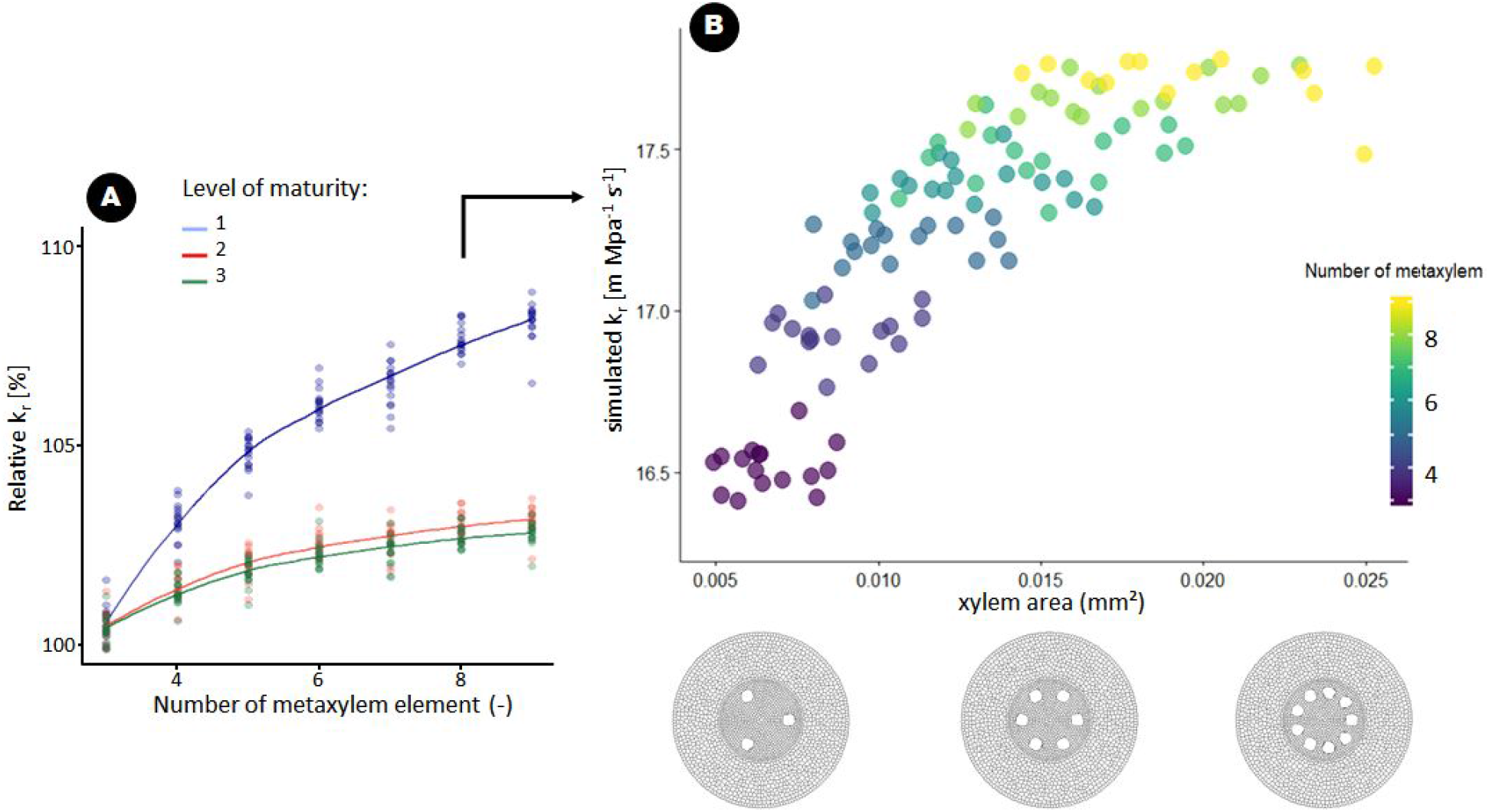
Influence of xylem features on the k_r_ for simulated monocot anatomies. **(A)** Influence of the number of xylem poles on the simulated relative k_r_. The colors represent different levels of hydrophobic barrier development. 1: Endodermal Casparian strip (blue); 2: Endodermal suberization (red); 3: Endodermis full suberization and exodermal Casparian strip (green). **(B)** Influence of the xylem area on the simulated k_r_. The simulated k_r_ was calculated using an endodermal casparian strip as the only apoplastic barrier.

**Figure 11:**
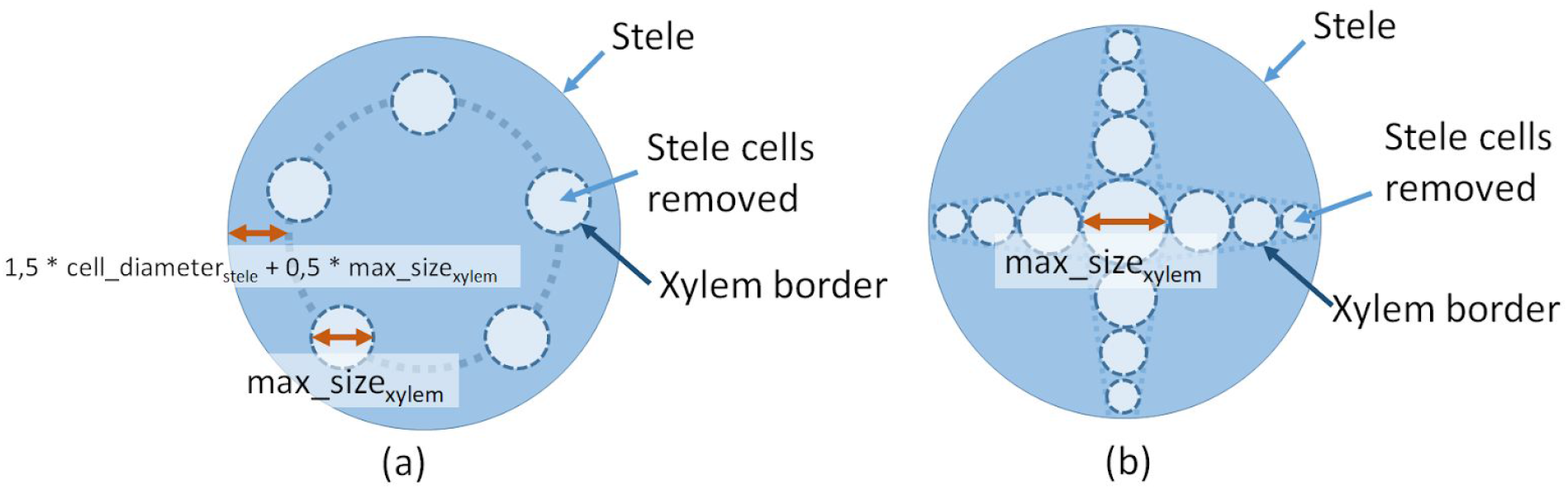
Overview of the methodology to create the vascular elements. (a) Monocotyledon stele cross section. (b) Dicotyledon stele cross section. In both (a) and (b), the stele cells inside the xylem border (dark blue) are removed to create xylem cells, indicated by light blue arrows. The different parameters used to build the root cross section simulation are described in light blue shaded boxes and indicated by double-headed red arrows.

## Discussion

### GRANAR streamlines the analysis of root cross sections anatomies

Previously, when analysing root anatomical data, scientists had to make compromises. On the one hand, a complete digitalization of the root anatomical network, that might be needed for further computational analysis, was possible using semi-automated tools, such as CellSet ^19^. However, this procedure is time consuming, especially for large roots. On the other hand, tissue-scale metrics could be automatically obtained from large datasets using other tools, such as RootScan ^21^, PHIV-RootCell ^22^ or RootAnalyzer ^23^. Such a high throughput pipeline yields quantitative descriptive properties, but they do not reach the level of details required for analyses of the functional role of root anatomy in functional structural plant models (FSPM)^2^

With GRANAR, we aimed to bridge the gap between these two pipelines, so researchers do not have to make tradeoffs between time consumption and quality of the analysis. Our new method uses tools that gather anatomical features to create GRANAR input parameters (examples are given in the Material and Methods). GRANAR then generates a complete root anatomical network which can be used for downstream root functional analysis (Fig. 2).

### GRANAR coupling with other tools enables the estimation of functional properties from root anatomical network

For instance, combining GRANAR with MECHA ^7^ enables the estimation of the k_r_ of a root cross section (fig. 2). Other computational tools, such as OpenAlea ^36^, could use the complete cell network anatomy generated by GRANAR, to analyse other functional properties, such as root radial growth. As such, cell-wall geometrical information can be integrated with mathematics to simulate how cell-tissue behave with different growth rates, local strain, and elastic deformation of cells under turgor pressure ^37^. For Arabidopsis thaliana, a single stereotypical root cross section was modeled by ^38^ to understand how cell properties and shapes contribute to tissue-level extensibility and yield. Their study could be extended to more diverse and bigger roots with GRANAR.

### Anatomical features largely influence the root radial conductivity

Overall, our simulations showed that the cortex width is one of the major predictors of k_r_. When the size of the cortex increases, the conductivity decreases and it is mostly due to an increase in the length of the apoplastic pathway, in our simulation cases. The larger the cortex is, the longest the path between the soil surface and the xylem vessels is. Even when apoplastic barriers are fully developed, the cortex thickness still influences the conductivity greatly (fig. 8). The influence of the cortex width on k_r_ was already pointed out experimentally for a wide range of species ^39^. Cortex features have also been pointed out to be a factor of drought tolerance ^8,9,40^. Several authors have shown that increasing the cortex cell diameter would reduce metabolic cost of soil exploration and consequently improve drought tolerance ^8,9,40^. In that regards, our results showed that for the same amount of cell layer, increasing cortex cell size would also reduce the k_r_ and limits water uptake. Reduced k_r_ would constitute a water saving strategy for the plant ^40^. Additionally, a reduction of the number of cortical layers has been shown to also reduce metabolic cost and therefore improve drought tolerance by enabling deeper rooting depth and increased water uptake ^8^. Here, our results showed that a reduction of the number of cortical layer would increase k_r_ and effectively ease water uptake. Thus, as suggested previously, roots with a small number of cortex layers and large cortex cells should have a small metabolic cost ^41^. However, our results showed that this would have a compensatory effect on k_r_ because few cortical layers and large cortical cells have an opposite effect (fig. 8c - 8d). We suggest instead a combination of few but large cortical cell layers that should show deep rooting ability thanks to the low metabolic cost and a small value k_r_.

Our simulations show that an increase in the aerenchyma proportion leads to a decrease in root k_r_. This has also been shown experimentally in maize ^42^ and barley ^10^. The reduction of the number of parallel pathways due to the formation of aerenchyma is likely the cause of the observed reduction of k_r_ (fig: 7). Increasing the aerenchyma has been shown to reduces the root metabolic cost, which in turn enables deeper rooting depth and improve drought tolerance ^43^. It has to be noted that the aerenchyma proportion is an anatomical feature independent from the many other anatomicals traits ^44^. We acknowledge that we do not know if cell hydraulic properties change with the aerenchyma formation in nature. This discrepancy may explain why the experimentally observed k_r_ decrease ^42^ is stronger than what we estimated with our simulation.

Recent studies showed that a large number of metaxylem vessels with small diameters improves drought tolerance ^45,46^. A hypothesis to explain the drought tolerance mechanism of small xylem diameter is the reduction of the cavitation likelihood ^47^. Another hypothesis is the reduction of axial water fluxes ^48^ which correspond to the water saving strategy ^40^. In addition, our results showed an increase of the k_r_ as the number of xylem poles increases. Moreover, a positive correlation between the number of xylem vessels and the k_r_ has been observed in *Ferocactus acanthodes* and in *O. ficus-indica ^49^*. Nevertheless, our simulation also showed that the decreasing size of xylem vessels alone would also decrease the k_r_ (fig. 10) which would lead to a water saving strategy. Another study, showed a drought tolerant plant (*Sorghum*) did express few xylem vessels and large cortex ^50^, in this case, the two factor would limit k_r_.

The stele area, to our knowledge, has not been explored as a drought tolerant factor in the field. However, experiment on maize lateral roots showed a strong correlation between stele diameter and the number of xylem poles which was discussed above ^51^. In our study, we simulate the change of stele diameter without changing the number of xylem poles, and already the stele alone has a strong impact on k_r_ (fig. 9). The increase of the simulated k_r_ as the stele area increases is mostly due to the increase of the pathway area.

### The influence of anatomical features changes with the development of apoplastic barriers

As detailed in the sensitive analysis for each of the anatomical features, the relative influence of each feature is generally decreasing as the number and strength of apoplastic barriers increase. The reason for this decrease is that the relative influence to water flow of the apoplastic barriers in comparison to anatomical variation is increasingly stronger. This behaviour typically occurred with the simulation for the cortex features and with the one for the xylem. Rieger and Litvin (1999) ^39^ justifiably, showed that apoplastic barriers alone do not explain all change of kr. However, our simulation shows that relative influence of anatomical variation changes as the apoplastic barriers increase (fig. 7–10). This, to our knowledge, has not have been pointed out in the literature. Further investigation could shed the light on experimental data, where such trends are observed.

### Current limitations and future developments

As shown in this manuscript, GRANAR is able to reproduce the general anatomical features of varied experimental data. However, the anatomies produced by GRANAR remain a stereotypical representation of root cross sections. Specific features, such as uneven cell divisions (for instance due to the formation of a lateral root) in specific locations will not be represented using GRANAR. As a results, scientists might want to use manual image analysis tools to capture such specific anatomical features. At this stage, it is also worth noticing that GRANAR does not represent structures formed during secondary growth (secondary xylem and phloem).

Another current limitations of GRANAR is its static aspect. The model does not explicitly simulate the growth and development of the different cell layers. The root anatomy is the result of a generative algorithm and is fixed in time. One possible improvement to GRANAR would be to include a developmental module within the model, similar to the current root architectural models ^52–54^. This would allow the model to create root anatomies for different root ages in one simulation, enabling the estimation of the evolution of the root conductivity with the root maturation. The down side would be that such model would be more difficult to parametrize, making it more difficult to link to experimental datasets.

A third limitation of our current pipeline is the lack of information regarding some of the cell layers. Although tools such as RootScan ^21^ or PHIV-RootCell ^22^ can automatically extract some of the needed variables, they do not extract all of them. For instance, some assumptions were made in our research regarding the cell size in the epidermis, endodermis, pericycle, stele and exodermis (see Material and Methods for details). It would therefore be useful to have a concerted development for both the model and the image analysis tools in order to create a fully integrated pipeline.

Finally, most of the data presented here were obtained on maize. This choice was a practical one, as maize was the only plant for which we could find sufficiently large and detailed datasets. In the future, additional validation of GRANAR should be performed, for other species. This would be facilitated if experimental data (in this case the images) were more widely shared within the community (e.g. on figshare or Zenodo).

## Conclusions

GRANAR is a new computational tool that is able to generate a range of root anatomies, for both monocots and dicots. The input of the models can be rapidly acquired using existing image analysis tools. As output, GRANAR produces an explicit representation of the root anatomy, as a network of connected cells. This networks can serve as input for other models, such as MECHA, and be used to estimate functional properties (here radial hydraulic conductivities) of specific anatomies.

In this manuscript, we validated GRANAR by re-analysing large published datasets of maize root anatomies. GRANAR was capable to represent the large range of experimental root anatomies. In addition, by coupling GRANAR with MECHA, we were able to estimate the hydraulic properties of the different anatomies in the dataset. This analysis highlighted the importance of cortex width and stele area as major factors influencing the root radial conductivity.

We also used GRANAR-MECHA to further explore the theoretical link between anatomy and radial conductivity. By creating 1000’s of theoretical anatomies, we were able to assess the functional importance of individual anatomical properties such as the number of xylem vessels, the proportion of aerenchyma or the size of the stele. The analysis highlighted that most anatomical variables have an influence on the hydraulic properties of the root. Their relative importances were variable and function of the level of apoplastic barrier development. As a general rule, the influence of anatomical feature was smaller when apoplastic barriers (in the endodermis and exodermis) were present.

In conclusion, we presented here a new computational pipeline for the functional characterization of root anatomy. GRANAR is an open-source project available at http://granar.github.io.

## Model and data availability

GRANAR is developed as an open-source project, under a GNU General Public Licence v3.0. Different sets of code and data were produced for this research:

- Web application: https://plantmodelling.shinyapps.io/granar/
- R package: https://github.com/granar/granar [10.5281/zenodo.3068505]
- R Markdown example: https://github.com/granar/granar_examples and [10.5281/zenodo.3068671]

## Authors contributions

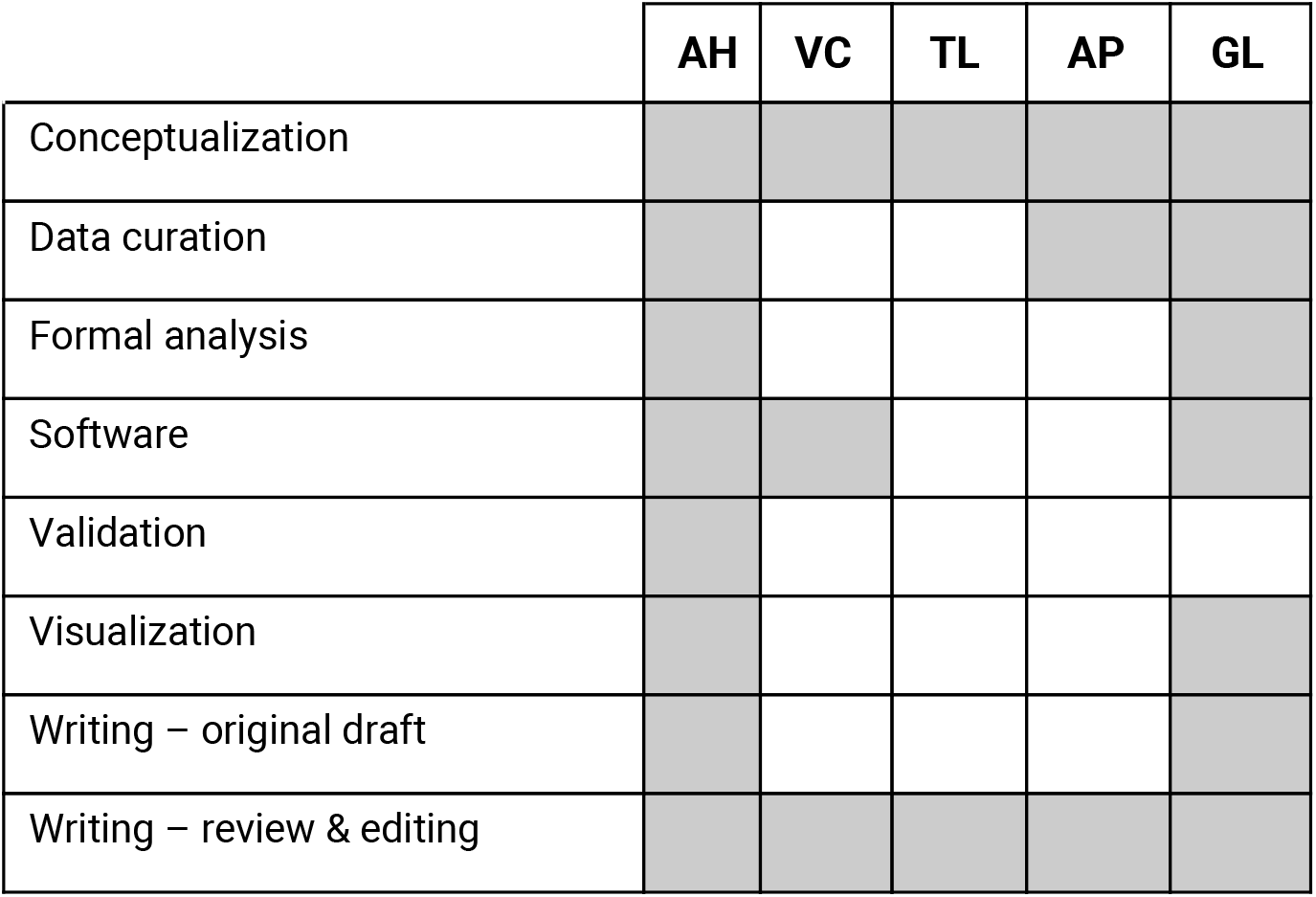

## Funding informations

This work was supported by the Fonds Spécial de Recherche (FSR) from UCLouvain (A.H.), the National Institutes of Health award number: T32GM007276 and NSF Graduate Research Fellowship (T.L.), the UCLouvain foreigners visitors program (T.L., A. P-G.), the Belgian National Fund for Scientific Research (FNRS, grant FC 84104) (V.C.), and the Communauté Française de Belgique-Actions de Recherches Concertées (grant ARC16/21-075) (V.C.).

## Material and methods

### Description of the GRANAR model

GRANAR was developed in R and is available as an R package [10.5281/zenodo.3068505].

To create a network anatomy of a root cross section, the GRANAR model is based on the placement of the cell layers around the center of the root. The model first places the innermost cell layer (the stele), then progressively builds the root toward the outermost layer (the epidermis). Coordinates of the cell center for each tissue type are established as a function of the diameter of the cell that compose the tissue. The cell type diameter is defined by the cell_diameter parameter for each cell type. The parameter n_layers defines the number of cell layers for each tissue type independently (Table 1). The only exception is for the stele, where the number of cell layers is calculated by dividing the diameter of the stele (layer_diameter) by the stele cell diameter. To create a unique network for every run of GRANAR, a random factor (randomness) add a slight variation to the coordinate of stele and cortex cell centers.

Specific vascular tissue patterns are created depending on whether the user choose to simulate a monocot or a dicot anatomy. For monocot, the xylem poles of a defined radius (max_size) are placed at the periphery of the stele. For dicots, the vascular blades are placed on defined radius from the center to the last layer of the stele. The size of the largest xylem element in the centrum is determined by the max_size parameter. The other xylem elements along the vascular blade have a decreasing size which is proportional to the distance from the centrum until it reaches the minimum size of a stele cell diameter. Once all cell centers are placed, the model creates the cell boundaries (cell walls). A voronoi tessellation of the cell centrums (using the deldir package ^55^) creates the cell walls at mid distance between every cell center previously set. The epidermis cells do not have any neighbour cells on the outer part, however an outside layer similar to the epidermis is created for the sake of the voronoi tessellation, then removed once it is done. Once the tessellation is done, the area of the different cells is then computed by the polygon function of the “sp” package.

### Description of the GRANAR package

The GRANAR model is available as an R package. The package contains the following functions:

`read_param_xml (path = “name_of_param_file.xml”)`

Load the parameter files that are under XML format and produce a dataframe object with the information needed to generate a root cross section with the function create_anatomy.

`create_anatomy (parameter = param_for_granar, path = “name_of_param_file. xml”)`

This function generates a root cross section with a data frame containing the needed parameter or calls the read_param_xml function to load a parameter file under XML format directly. The object returned by the function is a list containing four data frames:

- A `nodes` dataframe with the coordinates of all points of all the cells, their identity, and the information needed to create the cell polygons.
- A `walls` dataframe with the condensed information of all cell walls
- A `cells` data frame with the detailed information about the cells: area, distance from the center, and their angle in the cross section.
- An `output` dataframe with details informations for each tissue.

`get_root_section(path = “name_of_cross_section.xml”)`

The function imports the geometry information (the nodes dataframe) about root cross section from the file generated with GRANAR or CellSet.

`plot_anatomy(sim = simulation, col = “type”)`

Makes a figure which represents the given geometrical data. Accepted arguments for the col (color) parameter are `‘type’, ‘area’, ‘dist’, ‘id_cell’` and `‘angle’.` Default = `‘type’`

`write_anatomy_xml (sim = sim, path = “name_of_sim.xml”)`

Saves the geometry information generated with GRANAR in a XML file (the same output format as CellSet)

The package can be installed very easily on R thanks to the `devtools` package ^56^ with the function `install_github (“granar/granar”).`

The `granar` package rely on few dependencies. These dependencies are: the `deldir` package ^55^, the `alphahull` package ^57^ and the `retistruct` package ^58^. They should be installed (e.g. `install.packages(deldir)`) before loading the `granar` package.

### Description of the MECHA - GRANAR connection

MECHA reads XML-formatted CellSet data to load the geometry information that it need to compute the radial conductivity. GRANAR was build in order to generate outputs in the same format as CellSet. Therefore the coupling of those tools is easily made and examples can be found at this address: https://github.com/granar/granar_examples [10.5281/zenodo.3068671]. The coupling is done by file exchange.

### Description of the experimental data used for the validation of maize root anatomies

The Burton et al. (2013)^13^ data-set used in this study was the anatomical features of 195 landrace emphasizing accessions from stressful soil environments. The data is available in their supplement 1. The Chimungu et al. (2014)^9^ data-set used in this study was imported from their Supplemental Table S3 and Table S4 which is a list of genotype anatomical features selected from IBM population and another list of genotype anatomical features selected from Malawi maize breeding program.

The extraction of the anatomical features of Gao et al ()^27^ data-set was done with ImageJ software. The anatomical features gathered were exactly the same as the input parameter of GRANAR at the only exception of the used units. The anatomical features data used in our study can be found in their Table 1. The aerenchyma features of Gao et al. (2015) was found in their Figure 8. The Gao et al. (2015) data-set is the anatomical features of three root types of Zea mays L., cv. Zhengdan958 confronted to a low nitrogen treatment.

To create the three replicates of each data points, three run of GRANAR were made and saved separately for each data points.

### Description of MECHA hydraulic parameters

The simulation framework MECHA ^7^ estimates k_r_ from the root transverse anatomy generated with GRANAR or CellSet ^19^, and from the sub-cellular scale hydraulic properties of walls, membranes, and plasmodesmata. Cell wall hydraulic conductivity was set to 2.8 10^−9^ m^2^s^−1^MPa^−1^, as measured by ^59^ in maize. Lignified and suberized wall segments in the endodermis and exodermis (Fig. 6) were considered as hydrophobic and attributed null hydraulic conductivities. The protoplast permeability (L_pc_, 7.7 10^−7^ m s^−1^MPa^−1^) measured by ^60^ was partitioned into its three components: the plasma membrane intrinsic hydraulic conductivity (k_m_), the contribution of aquaporins to the plasma membrane hydraulic conductivity (k_AQP_) and the conductance of plasmodesmata per unit membrane surface (k_PD_). The latter parameter was estimated as 2.4 10^−7^ m s^−1^MPa^−1 7^ based on plasmodesmata frequency data from ^61^ and the plasmodesmata conductance estimated by ^62^. By blocking aquaporins with an acid-load treatment,^60^ measured a k_AQP_ of 5.0 10^−7^ m s^−1^MPa^−1^. The remaining value of k_m_ after subtraction of k_AQP_ and k_PD_ from L_pc_ was 0.3 10^−7^ m s^−1^MPa^−1^. Each value of k_m_, k_AQP_, k_PD_, and L_pc_ was set uniform across tissue types. For details on the computation of k_r_, see^7^.

### Data analysis

All statistical analysis were made using R. No distinction between the three data-sets was made to compute R squared and p values in the validation experiment. All ANOVAs were performed using the aov function from ‘stat’ package.

## Supplemental material

**Supplemental 1:**
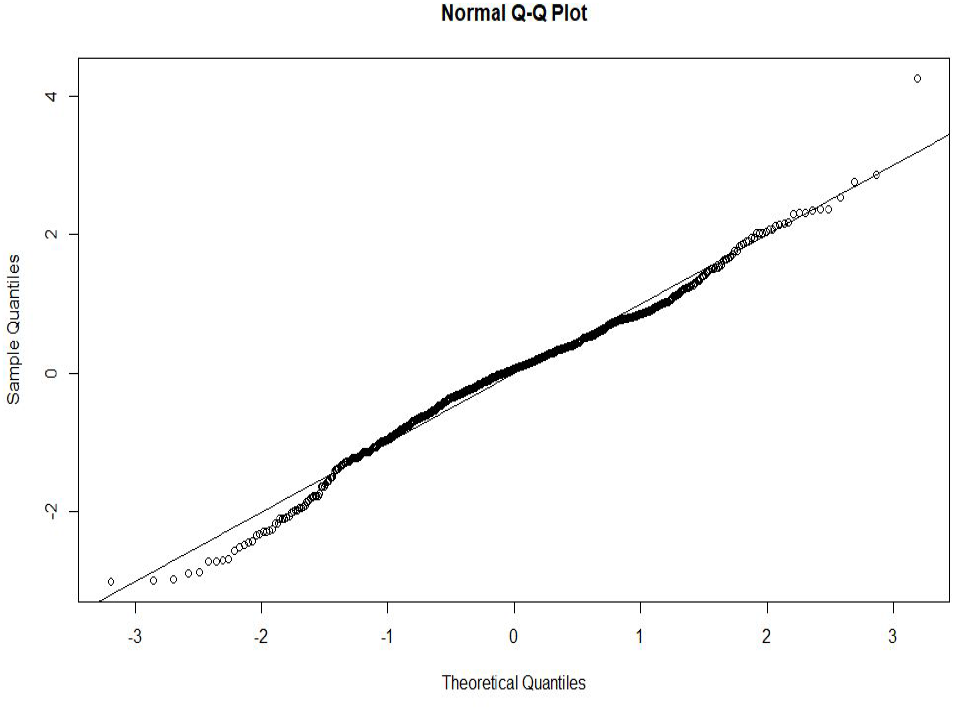
Quantile-Quantile (QQ) plot based on the residuals of the model summarised in equation 1.

**Supplemental data 1:** Spreadsheet containing the anatomical features and radial conductivity of all simulations. Tab “Fig. 3–5; Tab. 2–3” contains the data used and produced to re-analyse published datasets. Tab “Fig. 7” contains the data produced to do the sensitive analysis about the influence of the aerenchyma proportion on the k_r_. Tab “Fig. 8” contains the data produced to do the sensitive analysis about the influence of the different cortex features on the k_r_. Tab “Fig. 9” contains the data produced to do the sensitive analysis about the stele size on the k_r_. Tab “Fig. 10” contains the data produced to do the sensitive analysis about the influence of the xylem features on the k_r_.

